# Diet-dependent mortality and cognitive impairment reveal species-specific vulnerabilities to a microbial biopesticide in social bees

**DOI:** 10.64898/2026.04.17.718973

**Authors:** Flavia Di Cesare, Federico Cappa, Rita Cervo, Luca Ruiu, David Baracchi

**Affiliations:** Department of Biology, University of Florence, Via Madonna del Piano, 6, Sesto Fiorentino, 50019, Italy; Department of Agricultural Sciences, University of Sassari, Viale Italia 39/A, Sassari, 07100, Italy

**Keywords:** *Bacillus velezensis*, *Bacillus amyloliquefaciens*, *Apis mellifera*, *Bombus terrestris*, Nutritional stress, Synergistic effects, Risk-assessment, Sublethal effects, Learning and memory

## Abstract

The increasing use of microbial biopesticides in sustainable agriculture requires a deeper understanding of their potential impact on non-target pollinators. Although biocontrol agents are generally considered safer than synthetic pesticides, they may still cause subtle but ecologically relevant adverse effects on non-target organisms, especially when exposed to multiple stressors that are often overlooked in current risk assessment frameworks. Among these, nutritional stress, caused by habitat loss, fragmentation and reduced floral diversity, is becoming increasingly widespread. In this study, we investigated the lethal and sublethal effects of the bacterial biopesticide *Bacillus velezensis* (formerly *B. amyloliquefaciens*) strain QST713 at field-relevant concentrations on two key pollinators: *Apis mellifera* and *Bombus terrestris*. For the first time for a biopesticide, oral toxicity was assessed under environmental stress represented by diets with varying sugar concentrations (optimal and suboptimal) to identify potential synergistic effects on bee health. Sublethal effects were examined by studying learning performance and memory retention through a conditioning experiment under laboratory conditions. The results showed marked species-specific differences. While *B. velezensis* did not impact bee survival under realistic nutritional conditions, we observed a synergistic lethal effect in *B. terrestris* when biopesticide exposure was coupled with extreme nutritional stress (sugar deprivation). Similar species-specific differences emerged at the behavioral level: unlike *A. mellifera, B. terrestris* showed impaired visual learning and early long-term memory recall. Taken together, these results show that sublethal cognitive endpoints and multi-stressor contexts may reveal vulnerabilities not immediately evident through mortality-based assessments alone. Our findings also highlight the importance of including multiple pollinator species in risk assessment, as sensitivity to biopesticides might greatly vary among species.

## Introduction

Pollinators play a central role in both global food production and maintaining ecosystems. Over recent decades, populations of these species have declined due to multiple interacting stressors in agricultural and wild landscapes. The intensive use of synthetic pesticides has emerged as a major factor in pollinator losses (Cappa et al., 2022, 2025; Sánchez-Bayo & Wyckhuys, 2019). In response to these issues, biopesticides have emerged as a more sustainable alternative (Chandler et al., 2011). Biopesticides are pest control agents derived from natural sources such as microorganisms, biochemical compounds, or plant-incorporated protectants. They are being used more widely in organic agriculture and integrated pest-management because they are generally less persistent in the environment and considered less harmful to non-target species (Liu et al., 2021). The application of microorganisms as agents of biocontrol and biofertilizer has been recognized as an effective strategy to maintain crop productivity while reducing the application of chemical fertilizers and pesticides (De Andrade et al., 2023). However, the widespread assumption that biopesticides are inherently safe for pollinators and other non-target organisms is not consistently supported by empirical evidence. Current risk assessment frameworks are largely adapted from those originally designed for synthetic pesticides (Cappa et al., 2022). These protocols predominantly emphasize lethal endpoints, such as acute or chronic mortality, while sublethal effects, particularly those affecting behaviour, cognition, or physiological functions, are insufficiently explored. Even when considered, they are often limited to obvious symptoms such as reduced mobility or feeding activity. More subtle yet ecologically relevant alterations in learning, memory, or foraging efficiency are overlooked (Cappa et al., 2022; Mommaerts et al., 2009). Microbial biopesticides, such as *Bacillus velezensis* (formerly *B. amyloliquefaciens*), exemplify this knowledge gap. *B. velezensis* is a Gram-positive, spore-forming soil bacterium widely used in agriculture as a plant growth-promoting rhizobacterium and biocontrol agent (Qiao et al., 2014). It enhances plant growth and suppresses pathogens through the production of a wide range of bioactive compounds, including lipopeptides, polyketides, and lytic enzymes (Qin et al., 2015). Owing to these properties, it has been extensively applied in crop systems such as rice, cucumber, watermelon, and tobacco (Jiao et al., 2021; Saechow et al., 2018), especially as a fungicide, and is generally regarded as a sustainable alternative to synthetic pesticides (Luo et al., 2022). Although these microbial biopesticides generally exhibit low acute toxicity to bees, this apparent safety may mask biologically meaningful sublethal effects. For example, previous studies have shown that exposure to formulations containing *B. velezensis* spores can induce immunological alterations and reduce longevity in honey bees, despite negligible acute mortality (Sabo et al., 2020). Yet, behavioural endpoints, especially learning abilities and memory retention, remain unexplored in both honey bees and bumble bees following exposure to *B. velezensis* (Cappa et al., 2022; Cappa & Baracchi, 2024; Erler et al., 2022). These cognitive functions are fundamental for the ecological success of social bees, enabling foragers to memorize complex information about profitable floral resources and successfully navigate back to their nest (Baracchi, 2019; Raine & Chittka, 2008). Consequently, any cognitive impairment at the individual level can severely compromise foraging efficiency and the essential collection of resources, ultimately threatening the overall fitness of the colony (Klein et al., 2017).

Importantly, pollinators in real-world environments are rarely exposed to a single stressor in isolation (Vanbergen, 2013). Nutritional stress, driven by habitat loss and fragmentation, and reduced floral richness, is becoming increasingly pervasive. Limited or imbalanced food resources can compromise pollinator health and their resilience to other stressors (Woodard & Jha, 2017). Despite the long-standing recognition that interactions among stressors can amplify effects on biological traits, including those of biopesticides when combined with chemical agents (Isman & Norris, 2024), nutritional stress has been only marginally incorporated into pesticide risk assessments frameworks, and never examined in combination with biopesticides (Cappa et al., 2022). This omission is particularly concerning given that resource limitation represents a widespread condition in modern agricultural and urban landscapes. Consequently, the potential for interactive or synergistic effects between a promising microbial biopesticide such as *B. velezensis* and nutritional stress constitutes a critical, unexplored research frontier. Addressing this gap is therefore essential for developing more ecologically realistic and predictive risk assessments for non-target pollinators.

We evaluated both lethal and sub-lethal effects of *B. velezensis* on two key pollinator species: the honey bee, *Apis mellifera*, and the buff-tailed bumble bee, *Bombus terrestris*. Specifically, we (i) conducted mortality assays under nutritional-stress conditions and (ii) assessed associative learning and memory using conditioning paradigms, olfactory for honey bees and visual for bumble bees. By integrating both lethal and behavioural endpoints, our study provides a more comprehensive assessment of biopesticide impacts on non-target pollinators. Moreover, by adopting a comparative cross-species approach, we test whether responses observed in the honey bee can be generalized to other ecologically relevant pollinators such as bumble bees, thereby strengthening the ecological relevance of current risk assessment frameworks.

## Materials and Methods

### Insects

Experiments were performed on *A. mellifera ligustica* and *B. terrestris* workers. Honey bees were obtained from an experimental apiary consisting of five colonies located at the Department of Biology, University of Florence (Italy). Foragers returning to the hive were captured at the entrance and immediately transferred to the nearby laboratory.

Four bumble bee colonies were purchased from Koppert s.r.l. and maintained in laboratory conditions (23 ± 2 °C; 50 ± 10% RH; constant darkness) with *ad libitum* access to a 50% (w/w) sucrose solution and fresh pollen until testing. For all bioassays, bumble bees were collected directly from the nest. To ensure sample uniformity, medium-sized individuals were specifically selected, carefully avoiding those that were too large or too small.

### Biopesticide preparation

The study employed spores of *B. velezensis* strain QST713, recently reclassified from *B. amyloliquefaciens* subsp. *Plantarum*, and formerly identified as *B. subtilis* (Pandin et al., 2018), available in the microbial collection of the Department of Agricultural Sciences, University of Sassari (Italy), and originally derived from the plant protection product Serenade ASO (Bayer Crop Science). The pure spore suspension production system was implemented in the laboratory by growing the bacterium inside 1 L flasks with 200 mL LB broth maintained in an orbital incubator at 30 °C shaking at 120 rpm. The bacterial culture was carried out through successive inoculations, staring from a pre-inoculum initiated with heat-shocked spores. This pre-culture was grown to exponential phase and then transferred into a subsequent flask containing LB broth, where the culture was monitored and maintained up to 72 hours until achieving complete sporulation. Spore harvest was conducted by centrifugation for 20 minutes at 4 °C and 5,000 x g, followed by a purity check under a phase-contract microscope (Zeiss) before suspension in sterile water. Spore concentration was initially determined using a Neubauer chamber (Blaubrand), and throughout the study, the suspensions were periodically checked by colony forming unit (CFU) counts to ensure consistency. Stock concentrations of 1 × 10^9^ and 1 × 10^8^ spores/mL were prepared and stored at - 20°C until use in experiments with insects after appropriate dilution.

### Test suspensions

Honey bees and bumble bees were orally exposed for three consecutive days to one of three treatment suspensions, prior to the behavioural and mortality assays. The treatment consisted of a 50% (w/w) sucrose solution either alone (Control, C) or supplemented with *B. velezensis* QST 713 at 1 × 10^6^ CFU/mL (Low, L) or 1 × 10^7^ CFU/mL (High, H). Our selected concentrations (10^6^ and 10^7^ CFU/mL) of *B. velezensis* spores were derived from the application rate indicated on the commercial product (Serenade ASO) label and were intended to represent a realistic exposure scenario (10^7^ CFU/mL) and a lower one (10^6^ CFU/mL) for bees visiting treated vegetation (Bayer CropScience, 2020; European Food Safety Authority (EFSA) et al., 2021; Sabo et al., 2020). For *A. mellifera*, groups of 10-12 foragers were randomly assigned to Plexiglass cages (9 × 7 × 11 cm) equipped with 20 mL syringes providing the respective test suspensions. For *B. terrestris*, single workers were isolated in Petri dishes containing a pierced 1.5 mL Eppendorf tube filled with the test suspensions to allow feeding. All insects were kept in the dark at 23 ± 2 °C and 50 ± 10% RH.

### Survival assay under different nutritional conditions

In order to assess the potential interactions between biopesticide exposure and nutritional stress, individuals who had previously undergone the three-day treatment were assigned to one of three post-exposure dietary regimes differing in sugar concentration: (i) an optimal diet (Z50) consisting of a 50% w/w sucrose solution; (ii) a suboptimal diet (Z20) consisting of a 20% w/w sucrose solution; and (iii) a non-nutritive diet (Z0) consisting of water only. Fresh solutions were provided every three days throughout the experiment. For *A. mellifera*, workers from five colonies were housed in cages containing 10-12 individuals each. For each treatment-diet combination, 10 cages were set up, resulting in a total of 30 cages per diet (see Table 1). For *B. terrestris*, workers were collected from two colonies. From each colony, 20 workers were assigned to each treatment within each diet (40 bees per treatment per diet). This resulted in a total of 120 exposed individuals per diet (Table 1). Bees that died during the initial exposure period were excluded from the post-exposure mortality dataset.

**Table 1.** Sample sizes used in mortality assays for *B. terrestris* and *A. mellifera* under different dietary regimes. Number of individuals tested in each treatment group across complete sugar deprivation (Z0), suboptimal diet (Z20), and optimal diet (Z50). C = control solution; L = low dose; H = high dose of *B. velezensis*.

| Diet | Treatment | N° Bumble bees | N° Honey bees |
| --- | --- | --- | --- |
| **Z0** | C | 38 | 111 |
|  | L | 39 | 104 |
|  | H | 40 | 115 |
| **Z20** | C | 40 | 125 |
|  | L | 39 | 107 |
|  | H | 39 | 122 |
| **Z50** | C | 38 | 114 |
|  | L | 33 | 111 |
|  | H | 32 | 118 |

All cages were maintained under the same controlled environmental conditions described above until the death of all individuals. Survival data were then used to quantify treatment-related mortality and to assess potential interactions between biopesticide exposure and nutritional stress.

### Behavioral assay: learning and memory test

#### Honey bees

Learning and memory performance in *A. mellifera* were assessed using the classical Pavlovian conditioning of the proboscis extension reflex (PER) (Bitterman et al., 1983; Matsumoto et al., 2012). A total of 373 foragers from 5 colonies (see Table S1) were tested, with 125 in the control group (C), 123 in the high-dose group (H), and 125 in the low-dose group (L). After the three-day exposure to the test suspensions, bees were cold-anesthetized for five minutes and individually harnessed in plastic tubes by securing the thorax with a small strip of tape and fixing the head with a drop of low-temperature melting wax, allowing free movement of antennae and mouthparts. After recovery (30 min), each bee received 3 µL of 30% sucrose solution to standardize hunger levels and maintained in the dark for 2 hours before the start of the conditioning assays (Carlesso et al., 2020).

We used a differential olfactory conditioning procedure where bees received five rewarded presentations (CS^+^) of one odorant paired with 30% sucrose (unconditioned stimulus, US) and five unrewarded presentations (CS^−^) of an alternative odorant presented alone. Odorants were either 1-hexanol or nonanal, balanced across individuals (i.e., for half of the bees 1-hexanol was the CS^+^ and nonanal the CS^−^, and vice versa for the remaining bees). Odor presentations were pseudorandomized with an inter-trial interval of 10 min. During reinforced trials, 3 s after odour onset, the antennae and proboscis were touched with a toothpick soaked in sucrose solution to elicit PER. Both antennae were stimulated to avoid side bias (Baracchi et al., 2018). The timing of the odour delivery was automatically controlled using an Arduino Uno microcontroller board (Carlesso et al., 2020).

Memory retention was tested 1 h (short-term memory), 24 h (early mid-term memory) and 72 h (long-term memory) after conditioning by presenting both CS^+^ and CS^−^ without reward (Baracchi et al., 2020).

#### Bumble bees

The associative learning and memory of *B. terrestris* workers was assessed using a free-moving proboscis extension reflex (FMPER) paradigm (Cappa et al., 2025; Muth et al., 2018). This approach allows the assessment of visual associative learning in freely moving individuals. A total of 110 workers from two colonies were tested (see Table S2), with 38 individuals in the control group (C); 37 in the high-concentration group (H); and 35 in the low-concentration group (L). After three days of exposure to the solutions, the bumble bees were placed individually in a rectangular Plexiglas tube (12 × 3.5 × 3.5 cm) with lateral openings for ventilation. Bumble bees were free to move and explore the tube, except for the last 4 cm (where the stimulus was presented during the test phase). To standardize motivation, the bees were food-deprived for one hour prior to testing.

During the learning phase, two colored cardboard strips (one light blue and one purple) were inserted through two holes (2 mm in diameter) at the end of the Plexiglas tube. The gate was then opened to allow the bumble bee to approach the strips of paper. One strip was soaked in a 50% (w/w) sucrose solution (reward, CS^+^) and the other in a saturated saline solution (punishment, CS^−^). To prevent positional and color biases, the rewarded color and the side on which it was presented were counterbalanced across individuals and trials, respectively. Each bee underwent ten consecutive learning trials with a 10 min inter trial interval. A choice was recorded when the bee first contacted one of the two stimuli. Choices of the CS^+^ were scored as correct, while choices of the CS^−^ were scored as incorrect. Learning performance was measured as the proportion of correct choices across trials. Short- and early long-term memory were tested at 1 h and 24 h after the final learning trial, respectively. For each test, bees were re-exposed to the same two visual stimulus without any reinforcement, and the first choice was recorded as a memory response. Between the two tests, bees were individually maintained in Petri dishes in the dark and provided with 50% sucrose solution *ad libitum*.

### Statistical analysis

A mixed effects Cox regression (Allison, 2014) was used to estimate the effect of treatment groups on mortality rate under different nutritional conditions. Time-to-event data were analysed separately for each dietary treatment (Z0, Z20, and Z50) using Cox proportional hazards models (R package *survival*). Biopesticide treatment (control, low concentration (L), high concentration (H)) was included as a fixed factor, with the control group set as the reference. Survival time (hours or day) and event status (death) were used as response variables. As all individuals died during the experimental period, no censoring was applied. Model significance was assessed using likelihood ratio tests, and effect sizes were expressed as hazard ratios (HR) with 95% confidence intervals (CI). Learning performance during differential olfactory or visual conditioning was analysed using generalized linear mixed models (GLMMs) with a binomial error distribution and logit link function, using the *glmer* and *emmeans* functions from the R packages *lmer4* and *emmeans* respectively (Bates et al., 2015; Lenth, 2023). The response variable was the binary proboscis extension response (PER; response/no response) to the conditioned stimuli (CS^+^ and CS^−^) across acquisition trials. Fixed effects included treatment (trt), trial sequence (sequence), CS type (CS^+^ or CS^−^), and their interactions. Random intercepts accounted for individual identity and colony to account for repeated measures and colony-level variability. Model selection was performed using likelihood ratio tests and comparisons of Akaike’s Information Criterion (AIC). The significance of fixed effects in the selected models was assessed using Type III Wald chi-square tests. Post-hoc comparisons between treatment groups and the control were conducted using Dunnett’s test. Acquisition score (ACQS), defined as the total number of correct responses to CS^+^ across conditioning trials, was analysed using Kruskal-Wallis test followed by Dunn’s post-hoc tests with Holm correction for multiple comparisons.

Memory retention was assessed at different post-conditioning time points (honey bees: 1 h, 24 h and 72 h; bumble bees: 1 h and 24 h). At each time point, differences among treatments in the proportion of individuals exhibiting a conditioned response were analysed using Pearson’s chi-squared tests. When significant effects were detected, pairwise comparisons of proportions were performed with Holm-adjusted p-values. All statistical analyses were conducted in R (version 4.4.1).

## Results

### Survival assay under different nutritional conditions

In honey bees, survival was not significantly affected by biopesticide exposure under any of the tested nutritional conditions (Figure 1). Specifically, no treatment effects were detected under severe nutritional stress (Z0, water only; likelihood ratio test: χ^2^ = 1.74, df = 2, *p* = 0.42), sub-optimal nutritional conditions (Z20; χ^2^ = 2.36, df = 2, *p* = 0.31), or optimal nutritional conditions (Z50; χ^2^ = 0.67, df = 2, *p* = 0.72) (see Table S3).

**Figure 1.**
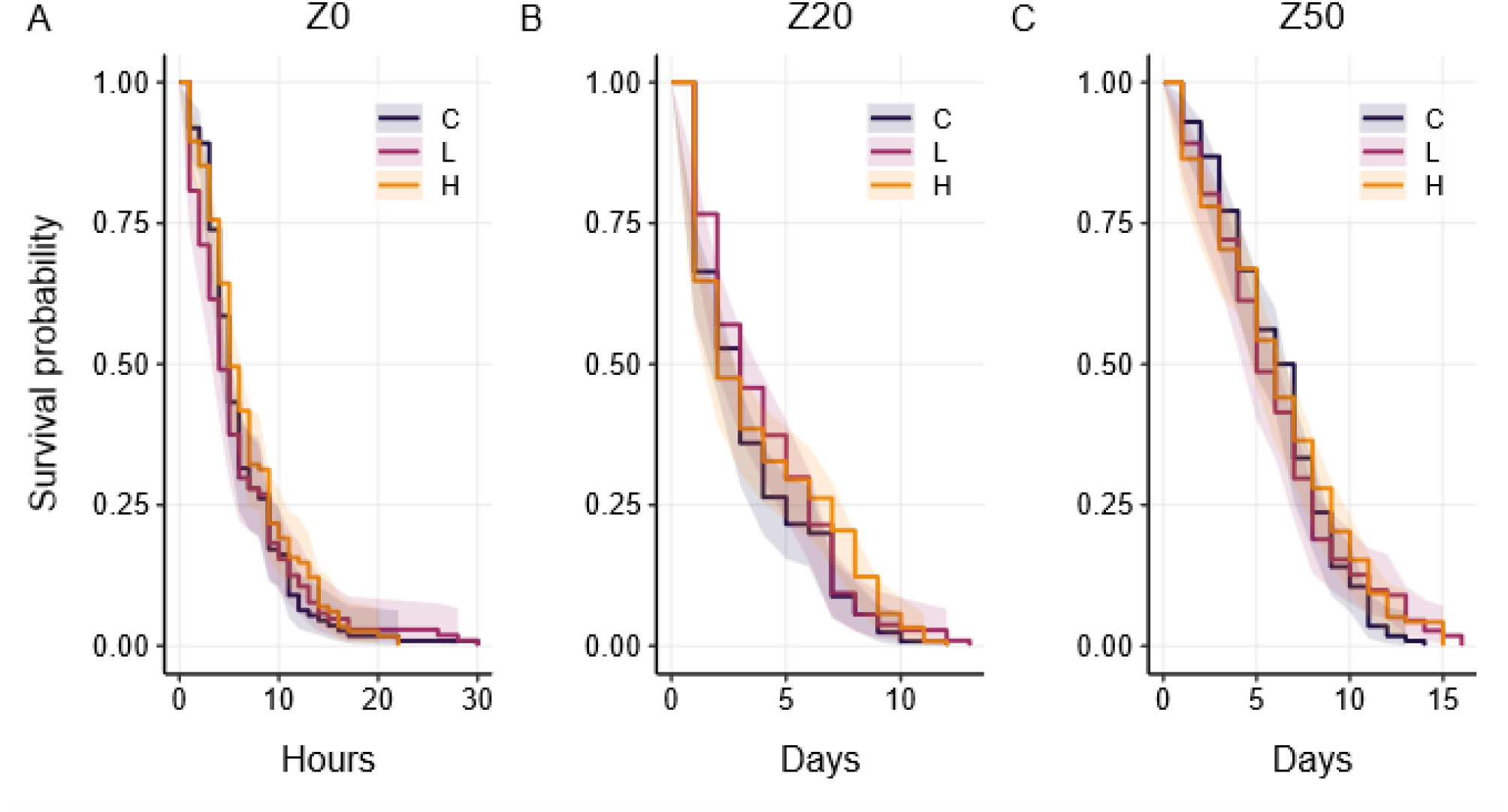
Effects of *B. velezensis* exposure on survival of honey bees under different dietary regimes. (A) Survival under complete sugar deprivation (Z0), (B) suboptimal diet (Z20), and (C) optimal diet (Z50) in honey bees exposed to control solution (C), low dose (L), or high dose (H) of the biopesticide. No significant treatment effects were detected under any dietary condition.

A similar pattern was observed in bumble bees under optimal and sub-optimal dietary regimes (Figure 2). Biopesticide treatment did not significantly affect mortality under the suboptimal Z20 diet (χ^2^ = 3.08, df = 2, *p* = 0.21) or under the optimal Z50 diet (χ^2^ = 2.91, df = 2, *p* = 0.23). In contrast, under conditions of complete sugar deprivation (Z0), the Cox model indicated a significant effect of treatment on survival (χ^2^ = 7.88, df = 2, *p* = 0.019) (Figure 2, A). Individuals exposed to the high biopesticide dose showed a significantly increased mortality risk compared to controls (hazard ratio > 1; Z0 High dose vs control: *p* = 0.005; Table S4).

**Figure 2.**
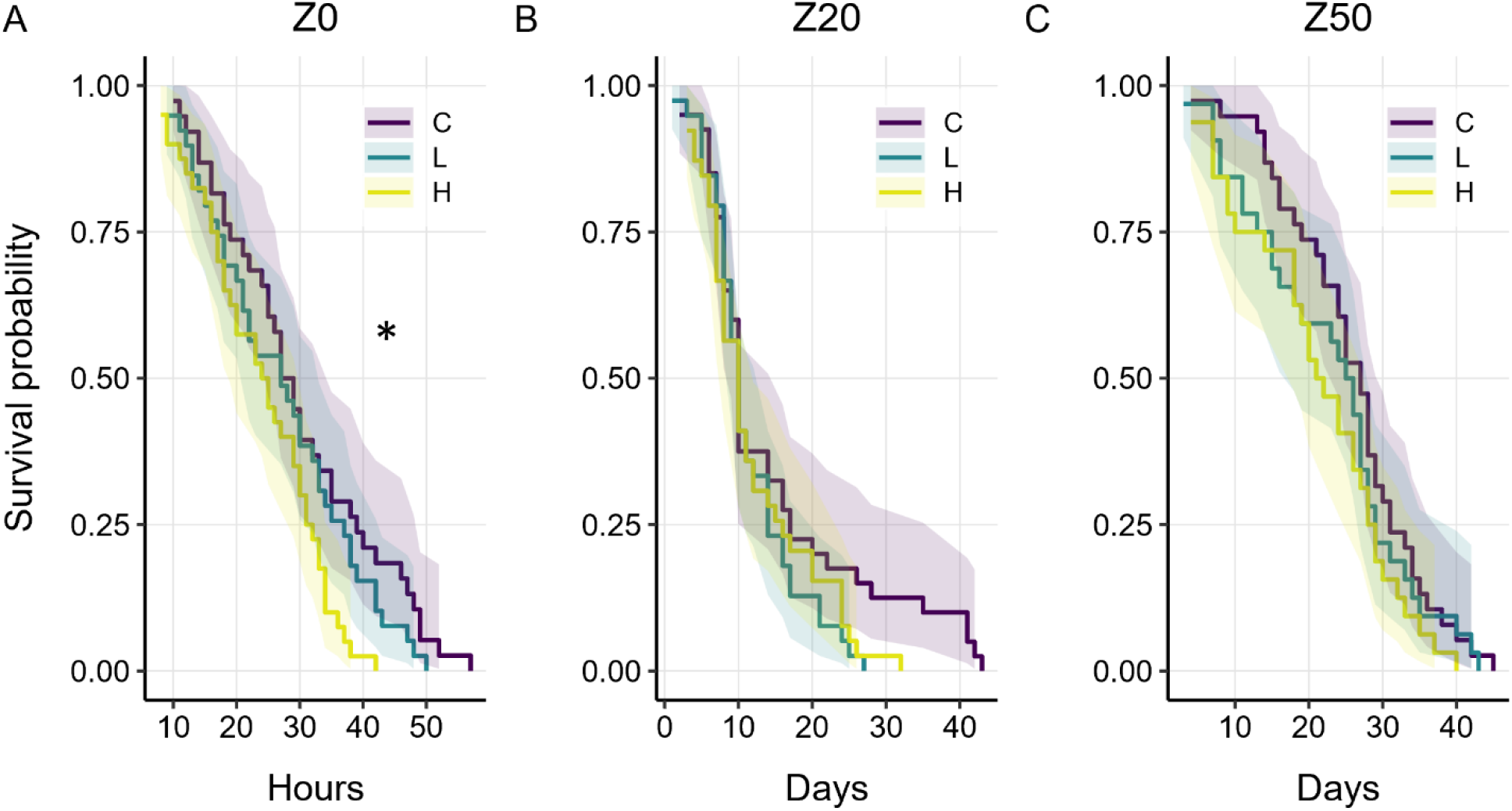
Effects of *B. velezensis* exposure on survival of bumble bees under different dietary regimes. (A) Survival rate under complete sugar deprivation (Z0), (B) suboptimal diet (Z20), and (C) optimal diet (Z50) in bumble bees exposed to control solution (C), low dose (L), or high dose (H) of the biopesticide. A significant treatment effect was observed under Z0 conditions (Cox model, *p = 0.019), with increased mortality at the high dose compared to controls, whereas no significant effects were detected under Z20 or Z50.

### Behavioral assay: Learning and memory test

#### Honey bees

During the acquisition phase of the olfactory conditioning, bees showed a significant increase in proboscis extension responses across trials (χ^2^ = 277.0, df = 1, p < 0.0001), and a clear discrimination between rewarded (CS^+^) and unrewarded (CS^−^) odors (χ^2^ = 357.1, df = 1, p < 0.0001). The biopesticide did not significantly affect learning performance (χ^2^ = 3.07, df = 2, p = 0.216), and no significant interactions with trial sequence or CS type were found (Figure 3).

**Figure 3.**
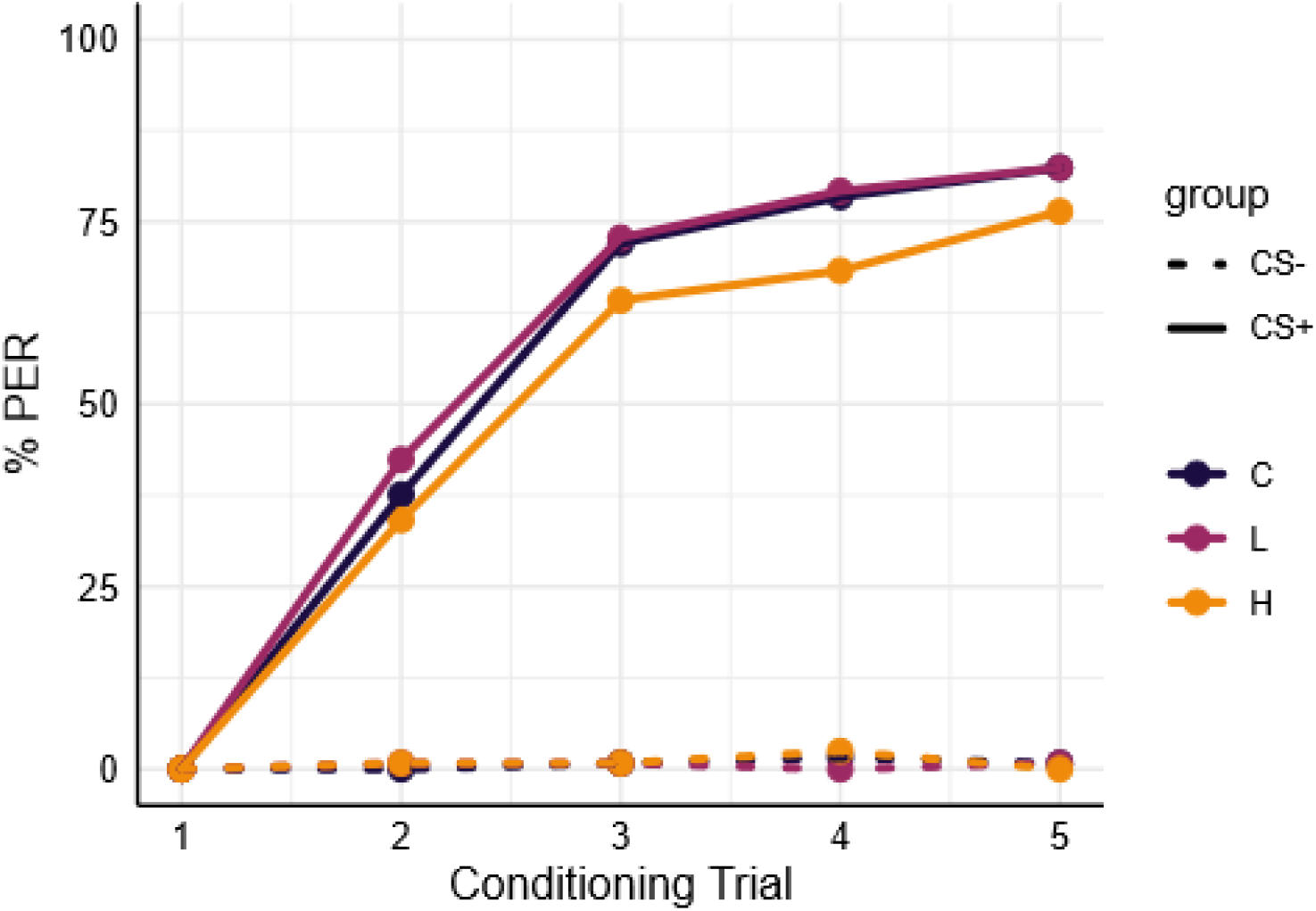
Effect of *B. velezensis* on associative learning in honey bees. Percentage of PER shown by bees during five conditioning trials when exposed to different concentrations (High (H), Low (L)) of biopesticides, or control (C) group. The solid lines represent the rewarded odorant (CS+) while the dotted lines represent the unrewarded odorant (CS-).

Memory retention tests conducted at 1 h, 24 h, and 72 h after conditioning revealed no significant differences in conditioned responses among treatments at any time point (1 h: χ^2^ = 0.115, df = 2, p = 0.944; 24 h: χ^2^ = 0.218, df = 2, p = 0.897; 72 h: χ^2^ = 0.769, df = 2, p = 0.681) (Figure 4).

**Figure 4.**
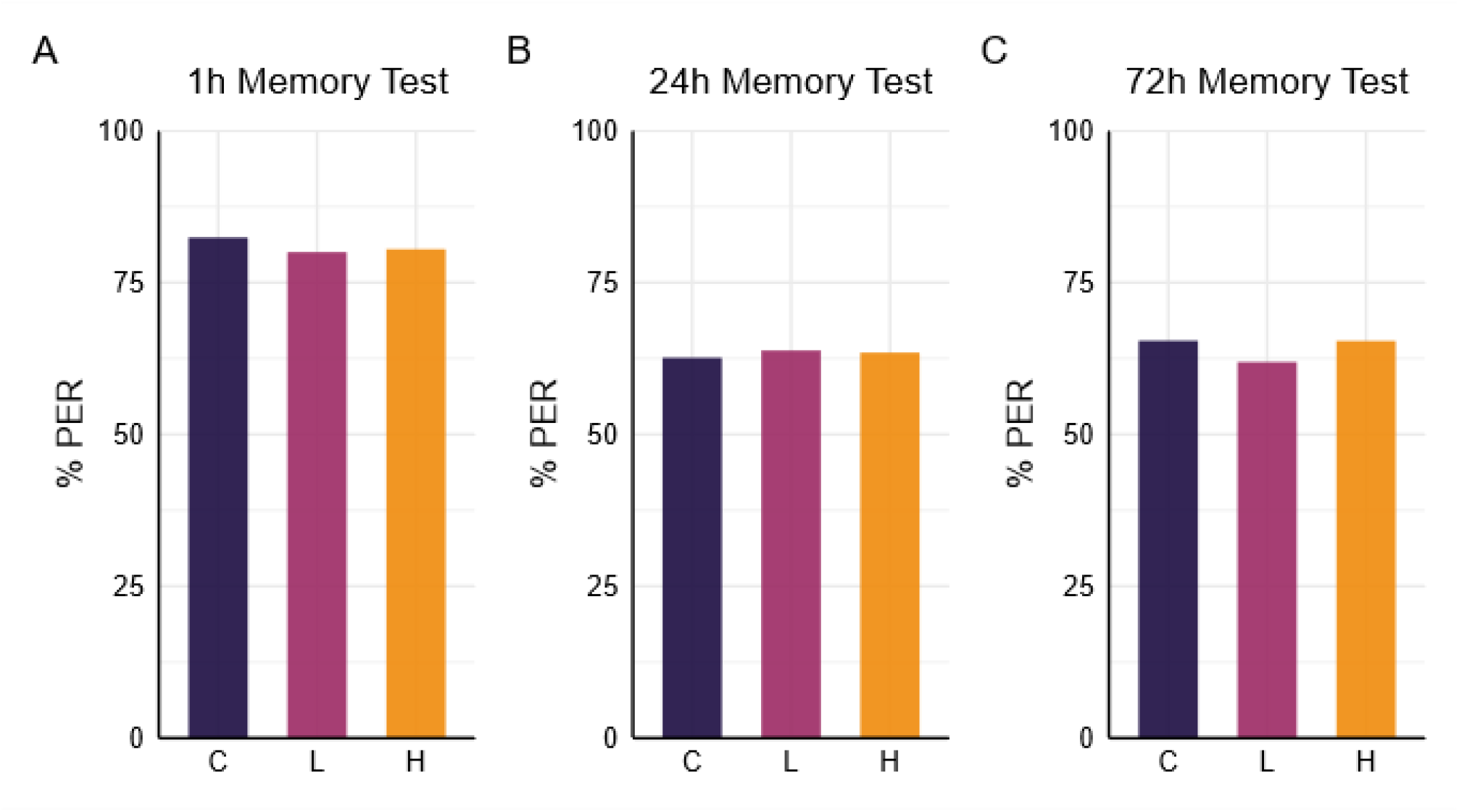
Effects of *B. velezensis* on memory retention in honey bees. (A) Short-term memory performance assessed 1 h after conditioning in bees exposed to control solution (C), low dose (L), or high dose (H) of the biopesticide. (B) Early long-term memory retention assessed 24 h after conditioning, and (C) long-term memory retention assessed 72 h after conditioning in the same treatment groups. No significant differences among treatments were detected at any time point.

#### Bumble bees

GLMM analysis of the learning phase revealed a significant main effect of trial 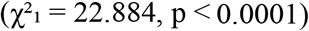, indicating an overall increase in the proportion of correct responses across trials. A significant main effect of biopesticides treatment was detected 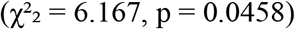. Post-hoc Dunnett comparisons showed that bumble bees exposed to the high-dose (H) of biopesticide exhibited significantly reduced learning compared to controls (z = -2.392, p = 0.0316), while the low-dose (L) group did not differ significantly from controls (p = 0.14) (Figure 5, A). Consistent with these results, the acquisition score (ACQS) differed significantly among treatments (Kruskal– Wallis: 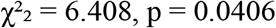). Dunn’s post-hoc comparisons revealed that the high-dose group achieved significantly lower acquisition scores than controls (adjusted p = 0.0470), while no significant difference was observed between the low-dose and control groups (adjusted p = 0.133) (Figure 5, B).

**Figure 5.**
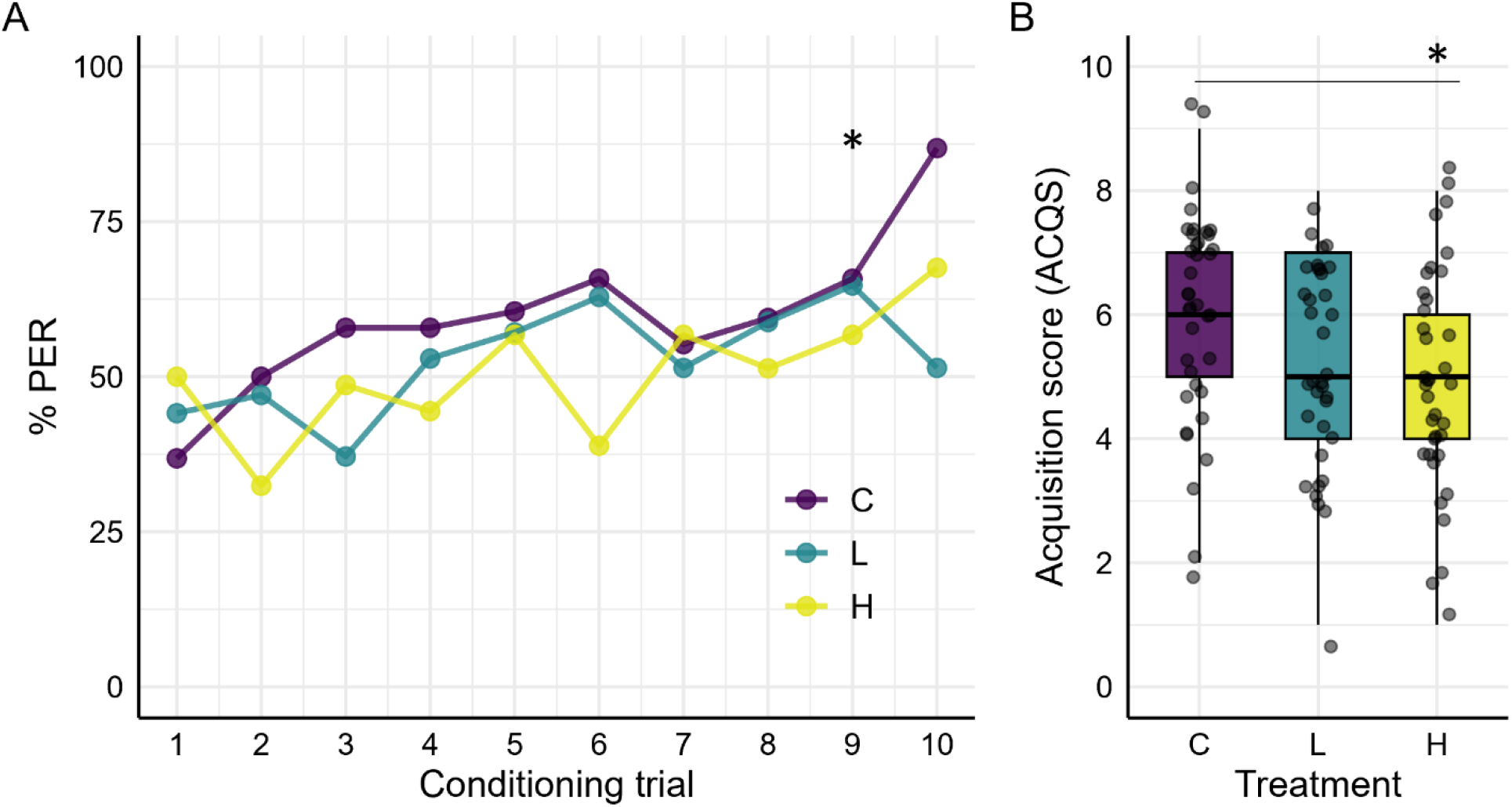
Effect of *B. velezensis* on learning performance in bumble bees. (A) Learning acquisition curves showing the proportion of correct responses across trials during conditioning trials in bumble bees exposed to control solution (C), low-dose (L) or high dose (H) of the biopesticide. Learning performance was reduced in the high-dose group compared to controls (*p = 0.0316), whereas the low dose had no effect. (B) Acquisition score summarizing overall learning performance across trials. Consistently, the high-dose group showed lower acquisition scores than controls (*p = 0.0470), while the low-dose group did not differ from controls.

Short-term memory performance at 1 h post-conditioning did not differ among treatment groups 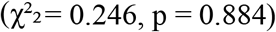 (Figure 6, A). In contrast, memory retention at 24 h post-conditioning differed significantly among treatments 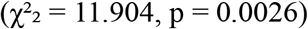 (Figure 6, B). Post-hoc pairwise comparisons of proportions (Holm-adjusted) revealed that both the high- and low-dose biopesticide groups showed significantly reduced memory retention compared to controls (control vs high: p = 0.0052; control vs low: p = 0.0148), whereas no significant difference was detected between the two biopesticide doses (high vs low: p = 0.8465).

**Figure 6.**
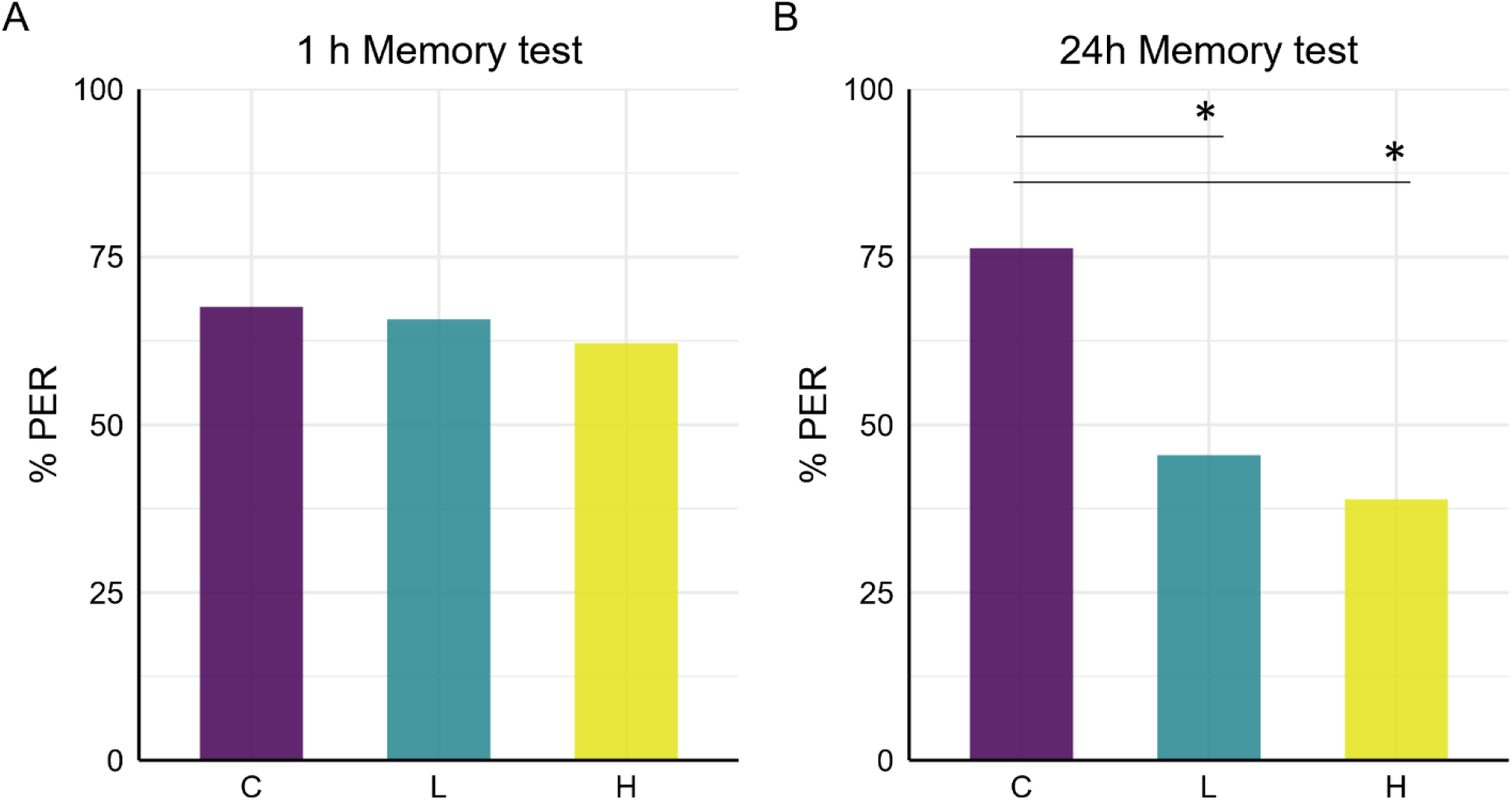
Effects of *B. velezensis* on memory retention in bumble bees. (A) Short-term memory performance assessed 1 h after conditioning exposed to control solution (C), low-dose (L) or high-dose (H) of the biopesticide. (B) Early long-term memory retention assessed 24 h after conditioning in the same treatment groups. No differences among treatments were detected for short-term memory (1 h). In contrast, at 24 h, both low- and high-dose groups showed significantly reduced memory retention compared to controls (*p = 0.0148 and *p = 0.0052, respectively), with no difference between biopesticide doses.

## Discussion

In this study, we investigated the lethal and sublethal effects of the bacterial biopesticide *Bacillus velezensis* QST713 on two key pollinator species, *A. mellifera* and *B. terrestris*. Survival was assessed under contrasting nutritional conditions, including a suboptimal diet (Z20) designed to reflect possible realistic scenarios of resource limitation, faced by pollinators in simplified agricultural landscapes or urban environments. In parallel, sublethal effects were evaluated by examining learning and memory performance. As previously highlighted, any disruption to these cognitive traits can severely impact individual foraging efficiency and homing ability, with cascading effects on colony provisioning (Klein et al., 2017). Overall, biopesticide exposure did not induce consistent lethal effects across the tested dietary realistic regimes. However, it does lead to species-specific sublethal impairments, with pronounced effects observed in bumble bees.

Under both realistic optimal and sub-optimal sugary dietary regimes, biopesticide exposure did not result in increased mortality in either honey bees or bumble bees, suggesting a limited lethal risk of *B. velezensis* QST713 at the individual level. These findings are consistent with previous laboratory studies reporting no lethal effects of *B. velezensis* on honey bees (Sabo et al., 2020) and bumble bees (Ramanaidu & Cutler, 2013). In contrast, Mommaerts et al. (2009) observed increased mortality in *B. terrestris* following a long-term chronic exposure, with most deaths occurring during the later phases of the experiment (after 11 weeks), highlighting the importance of exposure duration in shaping lethal outcomes. In natural landscapes, pollinators frequently encounter periods of nutritional limitation due to spatial and temporal fluctuations in floral resource availability, especially in areas heavily modified by agriculture or urban development, where floral diversity and nectar quality can be reduced (Woodard & Jha, 2017). Under such conditions, dietary stress may interact with other environmental pressures, including pesticide exposure, potentially amplifying their physiological costs. Therefore, assessing the effects of biopesticides under contrasting nutritional regimes provides a more ecologically relevant framework for understanding how multiple stressors can interact to influence pollinator health.

Under more extreme conditions, such as those imposed in the Z0 treatment, bumble bees, but not honey bees, exposed to the high biopesticide dose showed increased mortality compared with both the control group and the lower-dose treatment. Although complete sugar deprivation (Z0) does not represent a field-realistic feeding scenario at least for honey bees, which can typically rely on a large amount of stored food in the nest, it provides valuable physiological insight by acting as a stress-test that might reveal latent costs of exposure. Such costs associated with bacterial exposure may not necessarily translate into immediate mortality under field conditions but could nonetheless affect individual performance and ultimately colony fitness. This interpretation is coherent with previous studies reporting reduced reproductive output, such as decreased drone production in bumble bee microcolonies exposed to *B. velezensis* (Ramanaidu & Cutler, 2013). Why this effect was not evident in honey bees is difficult to explain with certainty, but it likely reflects species-specific differences in physiology, immune regulation, or metabolic resilience under nutritional stress. Although bumble bees are often reported to be less sensitive than honey bees to several stressor (e.g., microplastics and biopesticides) under standard laboratory conditions (Jütte et al., 2023; Pasquini et al., 2024), this pattern appears to be strongly context-dependent. In the present study, the increased mortality observed in *B. terrestris* in the Z0 groups may be partially explained by species-specific differences in starvation resistance. Individual bumble bees are generally more tolerant to food deprivation, as also showed by our results, than individual honey bees, surviving for longer periods under starvation. This prolonged survival could increase the temporal window during which energetic costs or pathogenic effects associated with bacterial exposure can manifest, allowing otherwise subtle effects to become detectable. Importantly, this interpretation does not necessarily imply that *B. velezensis* QST713 is inherently more detrimental to bumble bees than to honey bees. Rather, it suggests that differences in life-history traits and physiological resilience can modulate the expression of adverse effects. Similar considerations regarding the role of exposure duration are supported by Alkassab et al. (2024), who reported increased mortality in honey bees following chronic exposure to *B. velezensis* QST713, especially when applied in combination with other microbial control agents.

Cognitive performance measured under ongoing infections differed markedly between the two pollinator species. In honey bees, no significant impairments in learning acquisition or memory retention were detected following exposure to *B. velezensis* QST713, suggesting that *A. mellifera* may be relatively resilient to the cognitive effects of this biopesticide. In contrast, bumble bees exhibited a clear species-specific response, with high-dose biopesticide exposure impairing learning acquisition, and both tested doses negatively affecting early long-term memory retention. These results indicate that, although *B. terrestris* did not show consistent lethal effects under nutritionally realistic conditions, cognitive functions essential for efficient foraging might be particularly vulnerable to bacterial biopesticide exposure. Species-specific sensitivity to pesticides and biopesticides is increasingly recognised as a critical limitation of risk assessment frameworks largely based on the honey bee model (Cappa et al., 2022; Erler et al., 2022). Comparative studies have shown that different non-target species can exhibit contrasting lethal and sublethal responses to the same compound, even at field-realistic concentrations, reflecting differences in physiology, detoxification capacity, and life-history traits (Cappa et al., 2024; Jütte et al., 2023; Lima et al., 2024; Linguadoca et al., 2022; Lisi et al., 2025; Teixeira et al., 2022). Our results add to this growing body of evidence by demonstrating that sublethal cognitive endpoints may reveal vulnerabilities that are not detectable through mortality-based assessments alone. Together, these findings highlight that the emergence of adverse effects may depend not only on dose, but also on ecological context, underscoring the importance of assessing safety across a range of environmental conditions and in multiple non-target species providing crucial ecosystem services.

## Conclusion

Overall, our findings emphasize that bacterial biopesticides can induce subtle yet biologically relevant effects in non-target insects that are not captured by standard mortality-based assessments. The species-specific cognitive impairments observed in bumble bees, despite the absence of consistent lethal effects under realistic nutritional conditions, highlight the limitations of current risk assessment frameworks relying on a single model species, typically the honey bee. Cognitive assays targeting learning and memory represent functionally meaningful endpoints, as these abilities directly underpin critical behaviours such as efficient foraging, resource exploitation and navigation. Incorporating such sublethal endpoints into regulatory testing schemes would substantially improve the ecological relevance of non-target risk assessments.

While laboratory-based assays remain essential for isolating and standardising the effects of toxicants, our results also emphasise the need to integrate these approaches with semi-field and field-based studies. In particular, future studies should explicitly address potential consequences at the behavioural and ecological levels, including foraging efficiency and colony performance under realistic environmental conditions.

Importantly, our findings are not intended to diminish the importance of biopesticides, which represent a viable alternative to synthetic pesticides and generally have a lower environmental impact (Ayilara et al., 2023; Borges et al., 2021), especially in light of the growing need to support agricultural production. Rather, our results highlight the need for a more integrative framework for non-target risk assessment. This framework should combine multiple species, functional sublethal endpoints, and ecologically relevant exposure scenarios to better capture the complexity of pesticide-pollinator interactions across wild, agricultural, and urban environments.

## Acknowledgement

We thank Lucrezia Trucco and Bonacchi Martina for their valuable assistance with the experiments on honey bees and bumble bees.

