## Supplementary Material for "Diet-dependent mortality and cognitive impairment reveal species-specific vulnerabilities to a microbial biopesticide in social bees"

**Table S1: Sample sizes for the *Apis mellifera* learning and memory experiments.**

Number of honey bee workers tested per colony across the three experimental groups: control (C), high biopesticide dose (H), and low biopesticide dose (L), along with the total number of individuals per colony and per treatment.

| Colony | C | H | L | Total per colony |
|---|---|---|---|---|
| **1** | 24 | 23 | 22 | **69** |
| **2** | 30 | 29 | 28 | **87** |
| **3** | 25 | 22 | 27 | **74** |
| **4** | 25 | 25 | 25 | **75** |
| **5** | 21 | 24 | 23 | **68** |
| **Total per treatment** | **125** | **123** | **125** | **373** |

**Table S2: Sample sizes for the *Bombus terrestris* learning and memory experiments.**

Number of bumble bee workers tested per colony across the three experimental groups: control (C), high biopesticide dose (H), and low biopesticide dose (L), along with the total number of individuals per colony and per treatment.

| Colony | C | H | L | Total per colony |
|---|---|---|---|---|
| **1** | 19 | 20 | 18 | 57 |
| **2** | 19 | 17 | 17 | 53 |
| **Total per treatment** | **38** | **37** | **35** | **110** |

**Table S3: Effects of the biopesticide *B. velezensis* on honey bee mortality across different diets.**

Hazard ratios (HR), 95% confidence intervals (CI), and *p*-values were estimated using Cox proportional hazards models fitted separately for each dietary regimes: complete sugar deprivation (Z0), suboptimal diet (Z20), and optimal diet (Z50). Control honey bees (C) were used as the reference baseline (HR = 1.00) in all models. No significant treatment effects were detected under any of the tested dietary conditions ($p > 0.05$ for all comparisons).

| Diet | Treatment | Hazard ratio (HR) | 95% CI | p value |
|---|---|---|---|---|
| **Z0 (water only)** | L | 1.04 | 0.79–1.36 | 0.790 |
| **Z0 (water only)** | H | 0.88 | 0.67–1.14 | 0.325 |
| **Z20 (20% sugar)** | L | 0.84 | 0.65–1.09 | 0.197 |
| **Z20 (20% sugar)** | H | 0.84 | 0.65–1.08 | 0.171 |
| **Z50 (50% sugar)** | L | 0.95 | 0.73–1.24 | 0.702 |
| **Z50 (50% sugar)** | H | 0.90 | 0.69–1.16 | 0.413 |

**Table S4: Effects of the biopesticide *B. velezensis* on bumble bee mortality across different diets.**

Hazard ratios (HR), 95% confidence intervals (CI), and *p*-values were estimated using Cox proportional hazards models fitted separately for each dietary regimes: complete sugar deprivation (Z0, water only), suboptimal diet (Z20, 20% sugar solution), and optimal diet (Z50, 50% sugar solution). Control bumblebees (C) were used as the reference baseline (HR = 1.00) in all models. A significant treatment effect was observed under Z0 conditions (global $p = 0.019$), driven by increased mortality specifically at the high dose (H) compared to controls (HR = 1.97, $p = 0.005$), whereas no significant effects were detected under Z20 or Z50 conditions.

| Diet | Treatment | Hazard ratio (HR) | 95% CI | p value | |
|---|---|---|---|---|---|
| **Z0 (water only)** | L | 1.31 | 0.83–2.07 | 0.242 | |
| **Z0 (water only)** | H | 1.97 | 1.22–3.17 | 0.005 | *** |
| **Z20 (20% sugar)** | L | 1.46 | 0.91–2.33 | 0.113 | |
| **Z20 (20% sugar)** | H | 1.40 | 0.88–2.22 | 0.155 | |
| **Z50 (50% sugar)** | L | 1.21 | 0.75–1.94 | 0.435 | |
| **Z50 (50% sugar)** | H | 1.52 | 0.94–2.45 | 0.086 | |
